# Revealing The Colourful Side of Birds: Spatial Distribution of Conspicuous Plumage Colours on The Body of Australian Birds

**DOI:** 10.1101/647727

**Authors:** Kaspar Delhey

**Affiliations:** School of Biological Sciences, Monash University, Clayton, Victoria, Australia; Max Planck Institute for Ornithology, Vogelwarte Radolfzell, Radolfzell, Germany

## Abstract

In many species of birds, different body parts often display very different colours. This spatial distribution of coloured plumage patches may be determined by the balance between being cryptic to predators, and conspicuous to intended receivers. If this is the case, ventral and anterior body parts in birds–which are less visible to predators but more prominent to conspecifics– should present more conspicuous and sexually dichromatic plumage colours. Here I test these predictions using reflectance spectrometric measurements of standardised plumage patches across males and females for nearly an entire avifauna (Australian landbirds, N = 538 species). My data show that, as predicted, conspicuous colours are mainly located near the head, while the plumage of the back is the most cryptic. However, there is considerable variation across species, and this makes position on the body a relatively modest predictor of plumage elaboration (R^2^ = 0.15-0.19). One clear exception to this pattern is the conspicuous rump coloration. In many species, this patch can be concealed by wings, and therefore exposed only when necessary. In addition, conspicuous rump coloration could deflect or confuse predators in case of attack. Finally, patterns for sexual dichromatism were much weaker (R^2^ = 0.02), whereby wing and tail showed lower levels of dichromatism than the rest of the body.

## INTRODUCTION

Birds display a large variety of colours (Stoddard & Prum 2011) which are produced by several different mechanisms (Delhey 2015). This colour variation is partitioned between species (Gomez & Théry 2004; Dale *et al.* 2015; Delhey 2015), within species between populations, sexes (Dale *et al.* 2015; Delhey & Peters 2017) and also within an individual, since different parts of the body can have different colours (Fitzpatrick 1998; Friedman & Remeš 2015). This variation in colour between different body parts raises the question on whether this spatial distribution shows consistent patterns, with some colours being more prevalent in some parts of the body than in others.

Possibly the most pervasive and well-documented pattern of within-body variation is that darker colours are found dorsally and lighter colours ventrally on the body of a bird. This is known as countershading, and applies broadly to other animals as well (Cuthill *et al.* 2016). The main explanation for this pattern is camouflage, as countershading obscures the three-dimensional shape of the animal and improves background-matching (Cuthill *et al.* 2016). Other features of colour variation are however, less well documented. Nevertheless, balancing the trade-off between avoiding to be detected by unintended receivers (e.g. predators) versus being seen by intended receivers (e.g. potential mates) should lead to predictable patterns of colour distribution on the body of birds.

Cryptic colours should be more frequent in body parts that are most exposed to predators, and similarly conspicuous colours should be found where they are more visible to intended receivers. For most birds this would mean that cryptic colours should be found more often in dorsal regions which are more exposed to aerial predators (Gomez & Théry 2007). In addition, conspicuous colours, that play a prominent role in visual communication (Dale *et al.* 2015), should be more common in the anterior half of the body (head, breast, neck) because it is the more exposed region to intended receivers during face-to-face interactions (Doucet *et al.* 2007; Santana *et al.* 2012).

In general, support for these predictions comes from studies limited to few species or specific taxonomic groups (Stuart-Fox & Ord 2004; Doucet *et al.* 2007; Galeotti & Rubolini 2007; Gomez & Théry 2007). As far as I am aware, no systematic large-scale studies have assessed whether within-body spatial variation of colour conspicuousness follows the predictions outlined above. The only exception is Baker and Parker’s (Baker & Parker 1979) comprehensive analysis of the coloration of Western Palearctic birds, where they scored conspicuousness to human observers of several body parts but did not analyse within-body differences in a formal manner. Here, I use a large dataset comprising measurements of plumage reflectance across nearly all species of Australian landbirds (Delhey 2015) to test whether predictable differences in colour conspicuousness exist across body region of birds. I test these predictions by computing colour conspicuousness as the contrast between each plumage patch and common natural backgrounds. I also test whether there are consistent differences in sexual dichromatism across homologous patches in males and females (sexual dichromatism, (Delhey & Peters 2017)). Sexual dichromatism is thought to be a good indicator of sexual selection (Dale *et al.* 2015; Cooney *et al.* 2019), and I expect that this variable should follow similar patterns as outlined above. Hence, I predict that more conspicuous and sexually dichromatic colours should be found in ventral and anterior plumage patches.

## METHODS

### Reflectance measurements

Plumage reflectance spectra were obtained from museum specimens of 555 species of Australian landbirds as described in (Delhey 2015). On average 2.6 male and 2.4 female specimens were measured per species, and reflectance measurements were obtained from a set of 17 standardised plumage patches distributed across the entire body of the birds (Fig. 1). Bare body patches were not measured since these fade in coloration post-mortem and this is the reason why some species have fewer than 17 measured patches. For more details see (Delhey 2015). Reflectance measurements for both males and females were available for 538 species and all analyses were based on this subset.

**Figure 1.**
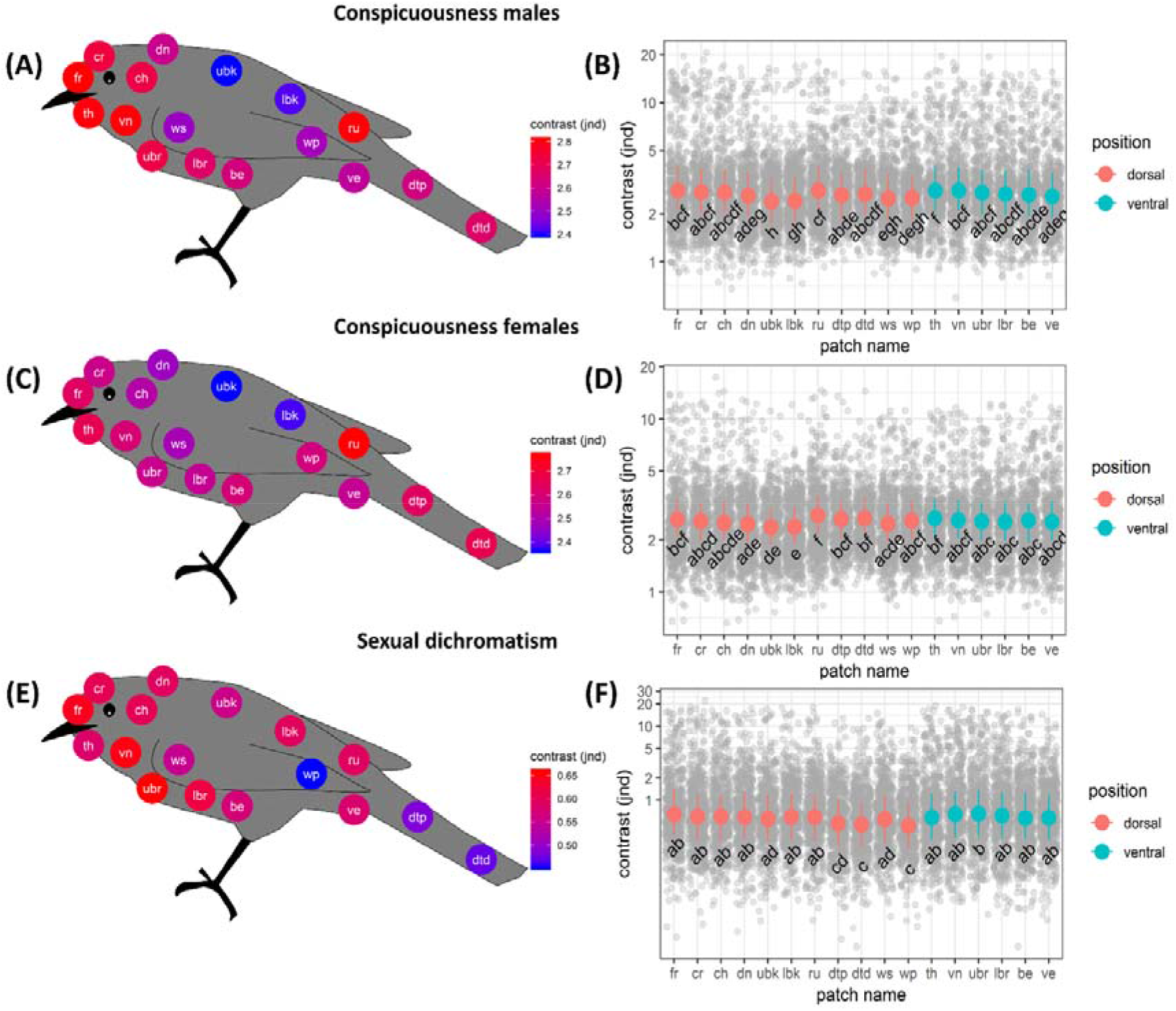
Spatial distribution of conspicuousness (chromatic contrast against natural backgrounds, A-D) and sexual dichromatism (chromatic contrasts between females and males, E-F) across 17 plumage patches on the body of Australian landbirds computed for birds with V-type visual system. Left-hand panels (A, C, E) show spatial distribution of plumage patches with colour gradient depicting mean contrast for males and females or sexual dichromatism; while right-hand panels shows model predicted means and 95% credible intervals depicted over raw data (grey symbols) for each plumage patch. Patches that share the same letter on graph have pairwise differences that are not statistically significant after Bonferroni correction (p < 0.00036). Effects and their 95% CIs are provided in Table S1. For a similar figure depicting effects computed using U-type visual sensitivities see Fig. S1. Patch abbreviations are as follows: fr = forehead, cr = crown, dn = dorsal neck, ubk = upper back, lbk = lower back, ru = rump, dtp = dorsal tail proximal, dtd = dorsal tail distal, ck = cheek, ws = wing covers, wp = wing primaries, th = throat, vn = ventral neck, ubr = upper breast, lbr = lower breast, be = belly, ve = vent.

### Visual models

Reflectance spectra (300-700 nm in 5 nm steps) were imported into the R environment and processed using psychophysical models of avian colour vision (Vorobyev & Osorio 1998). I used the formulas described in Cassey *et al.* (Cassey *et al.* 2008) as implemented by R scripts from (Delhey *et al.* 2015). These yield a set of three chromatic coordinates (xyz) that define the position of each reflectance spectrum in the visual space of birds. Euclidean distances in this space are measured in just noticeable differences (jnd) whereby distances of <1 jnd are unlikely to be perceivable by receivers.

Colour vision in most birds is mediated by four different types of single cones present in their retinas (Cuthill 2006), which are sensitive to very short (VS), short (S), medium (M) and long (L) wavelengths of light. Birds fall into two main groups based on their visual sensitivities: U- and V-type species, where the former have better UV sensitivity than the second (Hart & Hunt 2007; Delhey *et al.* 2013; Ödeen & Håstad 2013). I carried out the main analysis using V-type visual sensitivities since the main visual predator of birds are birds of prey, which have V-type visual sensitivity (Håstad *et al.* 2005), and the predictions I am testing concern the trade-off between avoiding being seen by predators and being seen by conspecifics. Nevertheless, I repeated all analyses using U-type visual sensitivities as well, to determine to what extent results are specific to one type of visual sensitivity. Visual sensitivities functions were obtained from (Endler & Mielke 2005). However, I emphasise most the results for V-type visual systems in the main text because most birds of prey have this type of visual sensitivity (Håstad *et al.* 2005), and therefore it is more directly linked to the risk of being detected by predators. As illuminant, I use the spectrum of standard daylight (d65, (Vorobyev *et al.* 1998) and noise-to-signal ratios for each cone type were computed using formula 10 from (Vorobyev *et al.* 1998), average cone proportions from (Hart 2001) (0.381:0.688:1.136:1) and a Weber fraction of 0.1 for the L cone (Olsson *et al.* 2017).

### Statistical analysis

Before the analyses, I averaged (separately for males and females) the chromatic coordinates of each plumage patch across the different specimens measured within species in order to obtain one set of xyz coordinates per patch for each sex and species (538 species, 9102 plumage patches for each, males and females). As an index of plumage patch conspicuousness I computed the contrast between each plumage patch and two common background types: brown bark and leaf litter and green leaves (backgrounds measured in Australia from (Delhey *et al.* 2013)). Background reflectance spectra were processed in the same way as plumage reflectance. To obtain a general index of conspicuousness I averaged the estimates of contrast against both background types (Delhey *et al.* 2017). While this does not capture the entire variety of background colours, it is largely representative of the rather limited palette of background colours present in terrestrial environments (Delhey *et al.* 2013; Cain *et al.* 2019). Indeed, values of conspicuousness were highly correlated (males: r = 0.97, p < 0.001; females: r =0.96, p < 0.001) with those obtained using reflectance spectra of natural backgrounds of European forests (Delhey *et al.* 2010). I also computed sexual dichromatism as the chromatic contrast between homologous plumage patches between males and females of the same species (Delhey & Peters 2017).

Response variables in the analyses were average chromatic contrast against the background (males and females separately), and sexual dichromatism. All response variables were log10 transformed to improve normality. I used Bayesian Phylogenetic Mixed Models (BPMM) as implemented in the R package MCMCglmm (Hadfield 2010). The only fixed effect was plumage patch (factor with 17 levels), while random effects included species identity (since each species contributes multiple plumage patches) and phylogenetic relatedness. Phylogenetic relatedness was accounted for by using the inverse of a phylogenetic covariance matrix. I incorporated phylogenetic uncertainty using the approach outlined in Ross *et al.* (Ross *et al.* 2013). Chiefly, a sample of 1300 phylogenies (based on the Ericson backbone) for the set of 538 species in the analysis was downloaded from www.birdtree.org (Jetz *et al.* 2012), and for each phylogeny, I ran the models over 1000 iterations saving the last sample. The latent variable values and variance components from this last iteration were used as a starting point for the next phylogeny. I iterated this over 1300 trees, discarding the results from the first 300 trees as burn-in, which resulted in a posterior sample of 1000 for each model. I used inverse gamma priors for the random effects and residuals, while for the fixed effects I used normal distributions centred on zero with large variances as priors. I evaluated model convergence by examining trace graphs and autocorrelation plots. I computed marginal (fixed effects) and conditional (fixed + random effects) R^2^ values, and also the separate contribution of each random factor (Nakagawa & Schielzeth 2010).

Data will be deposited in Dryad after acceptance.

## RESULTS & DISCUSSION

Based on V-type visual sensitivities, for both males and females, the most conspicuous colours were generally those situated ventrally and towards the head (Fig. 1A-D), and the least conspicuous plumage patches were located on the back. The combination of dorso-ventral difference in contrast against natural backgrounds, with less conspicuous dorsal regions, together with the already described countershading (Gomez & Théry 2007; Cuthill *et al.* 2016), which also occurs in this sample of Australian birds (Delhey 2018), probably contribute to make birds harder to detect by aerial predators such as birds of prey. There were some exceptions to this pattern however: the rump coloration showed higher levels of conspicuousness than other dorsal patches, and tail colours seemed also somewhat more conspicuous as well (Fig. 1). While standing in stark contrast to other dorsal patches in this analysis, a conspicuous rump is not a surprising sight to the seasoned birdwatcher: many bird species display distinctly coloured rumps, and my analysis confirms that this pattern applies broadly across an entire avifauna.

Why would the rump be highly conspicuous? I offer two, non-mutually exclusive, explanations. First, the tips of the wings often conceal the rump coloration when birds are at rest and so this plumage patch could be acting as a coverable badge (Hansen & Rohwert 2001) which can be exposed at will when needed as a signal. Second, rump coloration can also help evade predators after detection. For example, in feral pigeons (*Columba livia*) the contrasting rump coloration seems to distract the attention of attacking peregrine falcons (*Falco peregrinus*) helping them to escape (Palleroni *et al.* 2005). This would make rump coloration a deflecting mark (Caro & Allen 2017) or possibly constitute an example of dazzle or startle coloration (Caro *et al.* 2016) if the sudden flash of colour distracts an attacking predator when the prey takes flight. Conspicuous tail colours could also deflect attacks away from the body as is the case in lizards (Fresnillo *et al.* 2015).

Conspicuous colours seemed to be more frequently found towards the head (Fig. 1). Head colours would be most evident during face-to-face interactions such as with potential mates or rivals. This suggests that these body parts are more often involved in intraspecific communication (Dale *et al.* 2015). Indeed, complex face or head coloration patterns in primates and other mammals have been linked to sociality (Santana *et al.* 2012, 2013; Caro *et al.* 2017) and in manakins (Pipridae)–a polygynous clade of birds under strong sexual selection–male head colours were found to be most conspicuous (Doucet *et al.* 2007). Head coloration was also the more conspicuous body region across males and females of Western Palearctic birds (Baker & Parker 1979).

While differences in conspicuousness between plumage patches had statistical support, patch location on the body accounted only for moderate amounts of variation (marginal R^2^ males: posterior mean = 0.15, 95% credible interval (CI) = 0.127– 0.172; marginal R females: posterior mean = 0.19, 95% CI = 0.175–0.216) and differences in contrast across patches were relatively small relative to variation in the entire sample. Conditional R^2^ values, which include the contribution of fixed and random effects, were higher (males: posterior mean = 0.663, 95%CI = 0.614-0.713; females: posterior mean = 0.593, 95%CI = 0.549-0.633), mainly due to the strength of phylogenetic effects (phylogeny effect males: posterior mean = 0.513, 95%CI = 0.412–0.612; phylogeny effect females: posterior mean = 0.379, 95%CI = 0.281-0.472) since the contribution of species identity effects was smaller (males: posterior mean = 0.09, 95%CI = 0.0519–0.132; females: posterior mean = 0.114, 95%CI = 0.075-0.153).

Differences between plumage patches were even smaller for sexual dichromatism, which showed less clear patterns than conspicuousness (Fig. 1, marginal R^2^ posterior mean = 0.023, 95%CI = 0.019-0.027; conditional R^2^ posterior mean = 0.604, 95%CI = 0.541-0.672). The most sexually dichromatic patches seemed to be located towards the head and the least dichromatic on the wings and tail (Fig. 1), which again agrees with patterns described for manakins and Western Palearctic birds (Baker & Parker 1979; Doucet *et al.* 2007). Interestingly, rump coloration of Australian birds was not particularly sexually dichromatic (Fig. 1), which suggests that conspicuous rump coloration may be equally advantageous in males and females and therefore unlikely to be the result of strong sexual selection on males. In general, the weaker patterns for sexual dichromatism may result from the fact that most birds are sexually monochromatic, and even when dichromatic, sexes tend to resemble each other (Dale *et al.* 2015; Delhey 2015).

Patterns of variation based on conspicuousness estimates for U-type visual sensitivities showed similarities but also differences (Fig. S1, Table S1). While rump and head colours were still more conspicuous than the more cryptic back colours, lower ventral colours such as breast, belly or vent were generally less conspicuous. These differences highlight the fact that species with U-type visual sensitivities (mainly parrots, some gulls and some passerines (Ödeen *et al.* 2011; Ödeen & Håstad 2013)) may experience less stringent trade-offs between been seen by intended receivers and avoiding detection by birds of prey, which have V-type visual sensitivities, as has been suggested for European birds (Håstad *et al.* 2005).

In conclusion, analyses across an entire avifauna confirmed some of the predictions, even though the explanatory power of these models was rather low; the plumage patches presumably most exposed to predators show lower levels of conspicuousness than those in less exposed regions. Moreover, patches located on or near the head also showed higher levels of contrast. Since the rump seemed to be an exception to these patterns, we need behavioural studies to assess how rump coloration is used in interactions with conspecifics and particularly predators. While I suspect that the patterns described here are likely to apply to other regions of the world, more studies are needed to confirm this.

## DECLARATIONS

### Acknowledgements

I thank Jesse Smith, Hernan Morales and Roellen Little for help measuring museum specimens and the curators at the Melbourne Museum (Wayne Longmore, Karen Roberts) and the Australian National Wildlife Collection (Leo Joseph, Robert Palmer) for access to the specimens under their care.

