## Supplementary Material Fig S1 for "Revealing The Colourful Side of Birds: Spatial Distribution of Conspicuous Plumage Colours on The Body of Australian Birds"

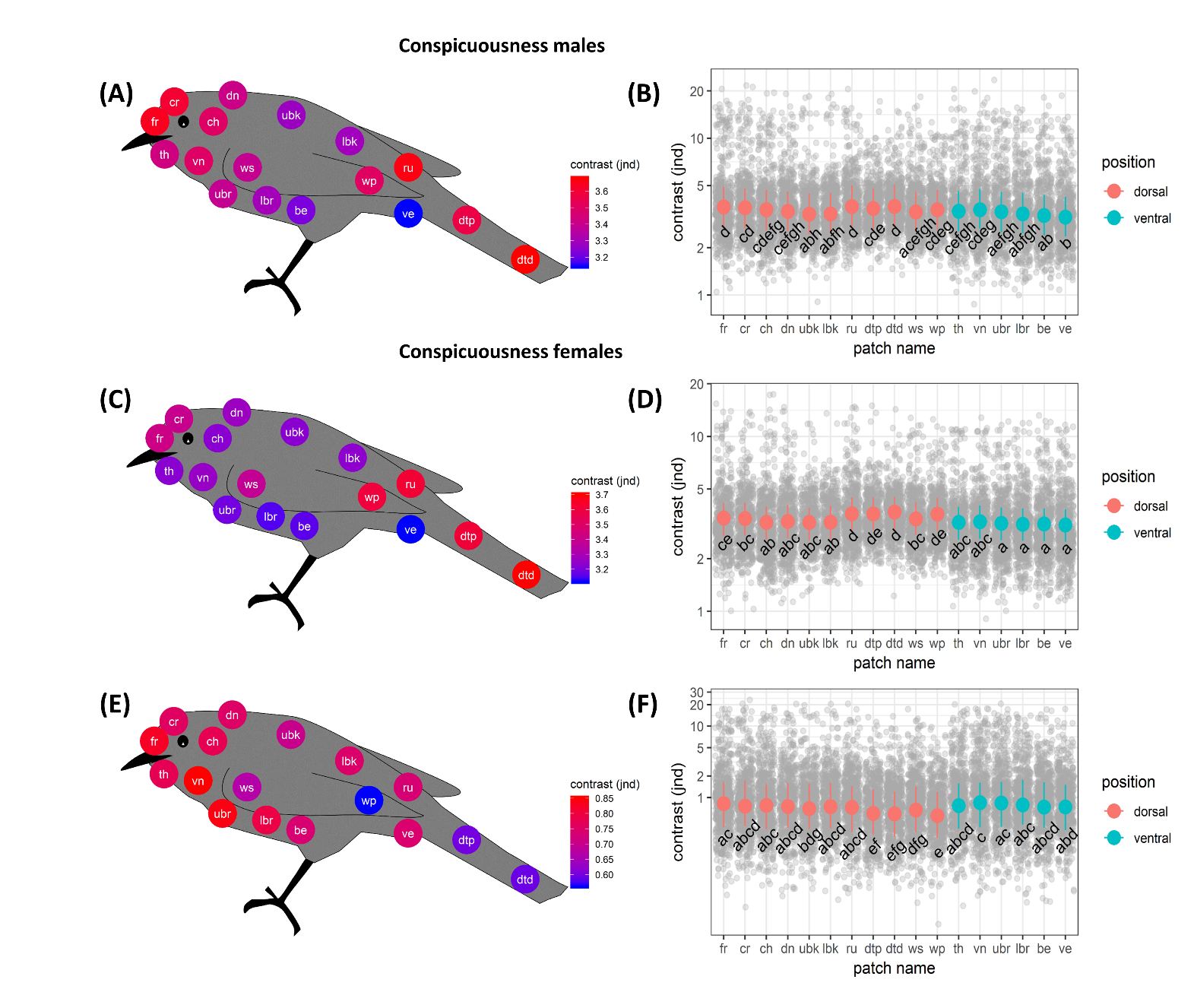

Fig. S1. Spatial distribution of conspicuousness (chromatic contrast against natural backgrounds, A-D) and sexual dichromatism (chromatic contrasts between females and males, E-F) across 17 plumage patches on the body of Australian landbirds computed for birds with U-type visual system. Left-hand panels (A, C, E) show spatial distribution of plumage patches with colour gradient depicting mean contrast for males and females or sexual dichromatism; while right-hand panels shows model predicted means and 95% credible intervals depicted over raw data (grey symbols) for each plumage patch. Patches that share the same letter on graph have pairwise differences that are not statistically significant after Bonferroni correction (p < 0.00036). Effects and their 95% CIs are provided in Table S1. Patch abbreviations are as follows: fr = forehead, cr = crown, dn = dorsal neck, ubk = upper back, lbk = lower back, ru = rump, dtp = dorsal tail proximal, dtd = dorsal tail distal, ck = cheek, ws = wing covers, wp = wing primaries, th = throat, vn = ventral neck, ubr = upper breast, lbr = lower breast, be = belly, ve = vent.

Table S1. Posterior means and their 95% credible intervals corresponding to values of male and female conspicuousness and sexual dichromatism for each of 17 plumage patches computed for V-type and U-type visual sensitivities as depicted in Fig. 1 and Fig. S1. Patch abbreviations are as follows: fr = forehead, cr = crown, dn = dorsal neck, ubk = upper back, lbk = lower back, ru = rump, dtp = dorsal tail proximal, dtd = dorsal tail distal, ck = cheek, ws = wing covers, wp = wing primaries, th = throat, vn = ventral neck, ubr = upper breast, lbr = lower breast, be = belly, ve = vent.

[table provided as separate csv file]
